# Accurate detection of HIV transmission clusters from phylogenetic trees using a multi-state birth-death model

**DOI:** 10.1101/215491

**Authors:** Joëlle Barido-Sottani, Tanja Stadler

## Abstract

HIV transmission networks are highly clustered, and accurate identification of these clusters is essential for effective targeting of public health interventions. This clustering affects the transmission dynamics of the HIV epidemic, which affects the pathogen phylogenies reconstructed from patient samples. We present a new method for identifying transmission clusters by detecting the changes in transmission rate provoked by the introduction of the epidemic into a new cluster. The method employs a multi-state birth-death (MSBD) model where each state represents a cluster. Transmission rates in each cluster decrease exponentially over time, simulating susceptible depletion in the cluster. This model is fitted to the pathogen phylogeny using a Maximum Likelihood approach. Using simulated datasets we show that the MSBD method is able to reliably infer both the cluster repartition and the transmission parameters from a pathogen phylogeny. In contrast to existing cutpoint-based methods for cluster identification, which are dependent on a parameter set by the user, the MSBD method is consistently reliable. It also performs better on phylogenies containing nested clusters. We present an application of our method to the inference of transmission clusters using sequences obtained from the Swiss HIV Cohort Study. The MSBD method is available as an R package.

## 1 Background

Basic epidemiologic models rest on the random mixing assumption (1; 2). In the presence of random mixing, each individual in a population has an equal probability of coming into contact with any other individual, which can lead to very quick epidemic spread. The random mixing assumption may be appropriate for airborne diseases in small communities. For sexually-transmitted infections (STIs) such as HIV-1 however, the random mixing hypothesis does not hold: STIs spread within sexual contact networks that limit their propagation to a specific subset of possible transmission events.

Identifying the structure in the sexual contact network has multiple applications, for instance allowing public health officials to target the populations most vulnerable to infection. One particular aim is to identify communities in the sexual contact network. These communities, or clusters, are defined as sets of nodes in the sexual contact network such that most or all nodes are connected within a cluster but few links exist between clusters (3). These clusters will affect the dynamics of an epidemic: at first the infection will spread quickly in the cluster where it has been introduced. The rate of transmission will then go down as the population of susceptibles in the cluster is progressively exhausted (2). Eventually a new introduction event may occur, where an individual from a previously uninfected contact cluster will be infected through one of the inter-cluster connections. Since the newly infected cluster is completely susceptible, the rate of transmission will then go up suddenly as new transmission routes open. Thus the cluster structure of the sexual contact network shapes transmission dynamics and thus may leave a detectable footprint in the phylogeny reconstructed from an epidemic. In what follows, we always consider phylogenies on the epidemic level, i.e. phylogenies obtained from pathogen genetic sequences of different infected individuals within an epidemic; thus each tip in the phylogeny represents a unique infected host.

Previous studies have found varying degrees of influence of the contact network on the phylogeny. (4) found almost no influence of the clustering coefficient of a network and the shape of transmission trees when the degree distribution of the network was kept constant, (5) found a modest effect of the degree distribution in the network on the shape of phylogenies reconstructed from simulated genetic data, whereas (6) found that the variance in degree distribution and the mean path length of the contact network could significantly affect the shapes of phylogenies. The link between network structures and phylogenies is also affected by viral characteristics such as within-host viral diversity (7). Several methods have been proposed to identify structural characteristics, such as connectivity and clustering coefficient, of the population network from a viral phylogeny (8; 9).

A number of methods have been proposed which exploit the effects that contact networks have on phylogenies to identify HIV transmission clusters from those phylogenies. In this paper we will focus on the methods evaluated in (10), which we will refer to as “cutpoint-based” methods. These methods differ in how they define the distance between two tips of the tree, but they have two major features in common: first, they require an ad hoc cutpoint to be specified by the user; second, they assume that the clusters are monophyletic in the phylogeny or monophyletic in a tree obtained from hierarchical clustering (Def. 4 in (10)), i.e that the most recent common ancestor of all tips belonging to a given cluster has no other descending tips. As (10) found, both features have a strong impact on the quality of the recovered clusters. Thus there is a need for a method which does not have these limitations.

Multi-state birth-death models have been widely used to model population structure and analyze phylogenies built from individuals in a structured population, both in epidemiological and macroevolutionary applications. Thus in principle such a model may be used to study the sexual contact network. In this context the aim is to infer which tips in a phylgeny belong to which cluster of an unknown contact network. Clusters differ by having different transmission dynamics through time, meaning different birth rates, so each cluster corresponds to a state in a multi-state birth-death model.

The Binary State Speciation and Extinction (BiSSE, (11)) and its extension to multiple states MuSSE, included in the package Diversitree (12), were the first efforts to infer state-specific birth and death rates from ultrametric phylogenies where each tip is assigned to a state. In (13), these approaches were extended to non-ultrametric trees. More recently the Beast2 package BDMM (14) allowed the joint reconstruction of a phylogeny and quantification of the parameters of an underlying multi-state birth-death model. These approaches require the user to specify how many states the model contains and to which state each tip of the phylogeny belongs. An exception to the latter is (13), which can integrate over tip states, but does not assign states to tips.

However we cannot readily use any of the above approaches to infer transmission clusters, for two reasons. First, the state of tips, i.e which cluster they belong to, is not known prior to the analysis. Second, integrating over the tip states instead explicitly assigning states to tips means that the repartition of tips into clusters cannot be inferred.

The method Bayesian Analysis of Macroevolutionary Mixtures (BAMM, (15)) addresses these issues and is able to infer the number of clusters and assign each tip to a cluster. Further, the birth- and death rate parameters associated with each cluster are quantified. However, it was designed to be used with macroevolutionary datasets, meaning at the time of this writing it can only analyze ultrametric trees, i.e with all tips sampled at the same point in time. For epidemiological datasets, we have non-ultrametric trees as samples are collected through time. Furthermore, its results have been called into question (16).

In this paper, we present a new method to identify clusters of transmission in a phylogeny built from viral sequences, by detecting ‘jumps’ in transmission rate. We associate these jumps with introduction events into previously untouched clusters. From the detected jumps, we can readily read off the partition of the tips of our phylogeny into distinct clusters. Our method uses the multi-state birth-death (MSBD) model with allowing decreasing transmission rates within clusters to account for the depletion of susceptibles. In particular, it does not require prior knowledge on the number of clusters or the tip repartition in clusters. We evaluate the performance of this new method on the simulated dataset of (10) and compare it to cutpoint-based methods. We then apply it to a published HIV phylogeny (9) which was obtained based on 192 sequences from the Swiss HIV Cohort Study. Finally we discuss the limitations of the method and planned future work.

## 2 Methods

## 2.1 Model

We use a multi-state birth-death model similar to the model used in the BDMM package (14). The birth-death process starts with one infected individual at time *τ* in the past in an ancestral state and is stopped at present time 0. This means that we measure time in the backwards direction, increasing from the present to the root. State changes happen in each individual through time with a rate *γ*. Our MSBD model contains an unknown number of states *n*,* corresponding to *n\** clusters in the underlying population network. We assume that all states are equally likely to transition to, so that the state change rate between any state *i* and *j* is as follows:

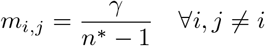

Each state *i* is characterized by a specific initial transmission rate *λ*_0,*i*_, a transmission decay rate *z*_*i*_, and a removal rate *M*_*i*_. Each individual produces an additional individual with a state-and time-dependent transmission rate *λ*_*i*_(*t*) (function of *λ*_0,*i*_, *z*_*i*_ as defined below), and is removed with a state-dependent removal rate *μ*_*i*_ corresponding to the rate of “becoming non-infectious”.

The depletion of the susceptible population is modeled by the exponential decay of the transmission rates in the process. Each state is associated with a specific initial transmission rate *λ*_0,*i*_ and a transmission decay rate *z*_*i*_ Equation 1 shows the transmission rate for a lineage in state *i* at time *t* before the present, where *t*_0,*i*_ is the time of introduction into state *i.* Since time is backwards, we impose *z*_i_ ≥ 0, so that the transmission rate decreases as the process progresses towards the present.

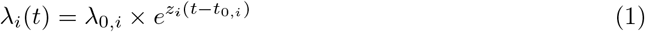

The infected individuals are sampled upon removal with a probability *σ.* This birth-death model produces a tree on all infected individuals together with position and times of rate changes on the tree, and we obtain the phylogeny by considering the subtree spanned by the sampled infected individuals. The phylogeny contains information about the transmission and removal times of the sampled individuals, as well as the positions and times of the rate changes, as shown in Figure 1. We assume that the state changes correspond to introduction events in newly infected clusters, so that all tips inferred to be in the same state belong to the same cluster in the original transmission network.

**Figure 1:**
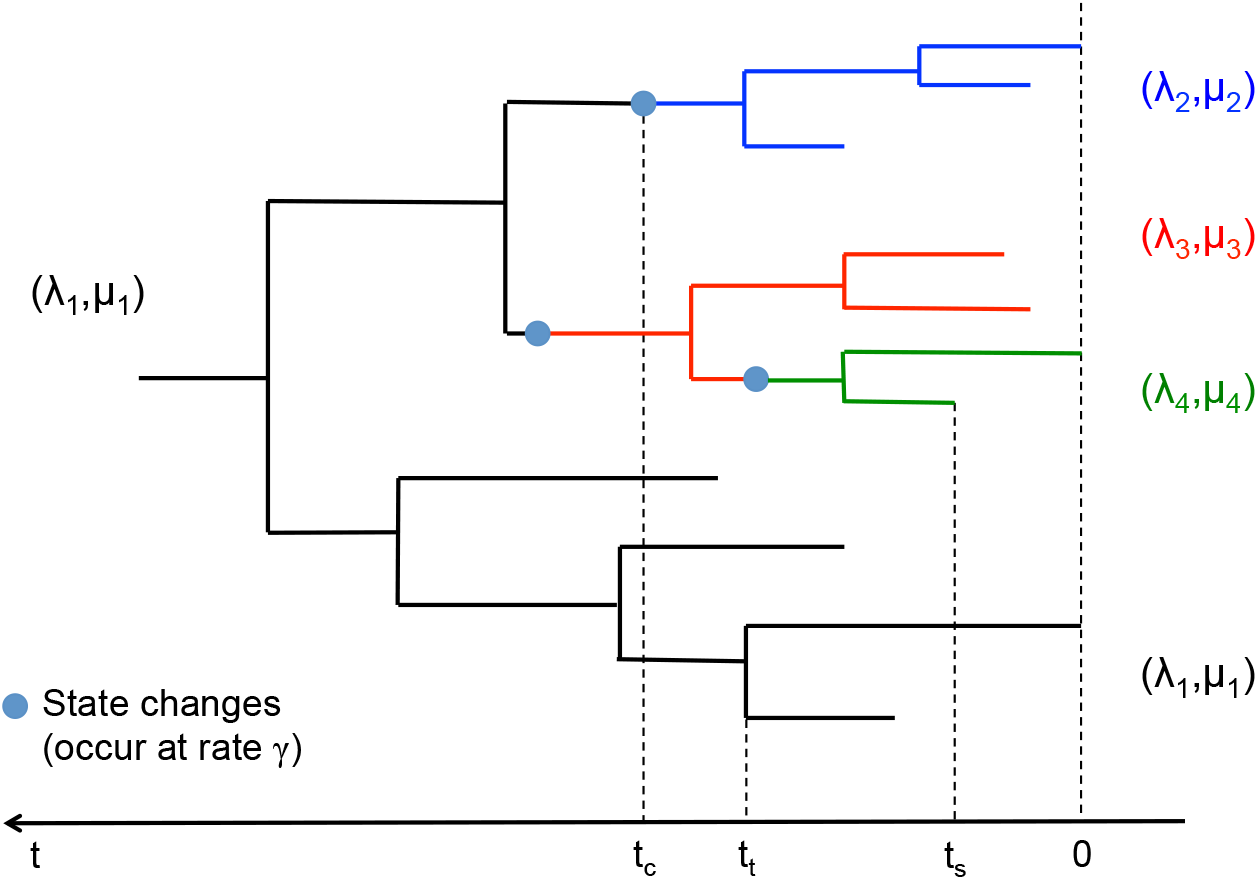
Visual representation of the phylogeny under a MSBD model. Each state is represented by a colour: the ancestral state, in black, starts at the root and represent the first cluster infected. The other states, in blue, red and green, start at change points along the tree. These states represent the clusters infected later in the course of the epidemic and the state change points represent the introduction event for each associated cluster.

We refer to a node in the phylogeny being either a branching event, a tip, or a state change event. Edges in the phylogeny connect any two nodes, and so any edge belongs to only one state.

## 2.2 Likelihood function

We now derive the probability density of a phylogeny (including the state change times) given the MSBD parameters, i.e. we derive the likelihood of the parameters given a phylogeny.

### 2.2.1 Differential equations

Following (13; 14), the likelihood function of the model parameters given the phylogeny can be calculated from the differential equations below. Eqn. (2) describes the probability *p*_*i*_(*t*) of a lineage in state *i* at time *t* not producing any sampled offspring until the present (referred to extinction probability below). Eqn. (3) describes the probability density *q*_*i*,*N*_(*t*) of an edge *N* in state *i* at time *t* evolving according to the phylogeny in time interval [*t*, 0].

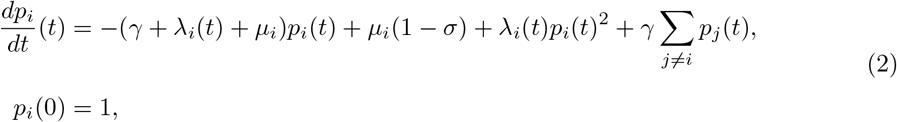

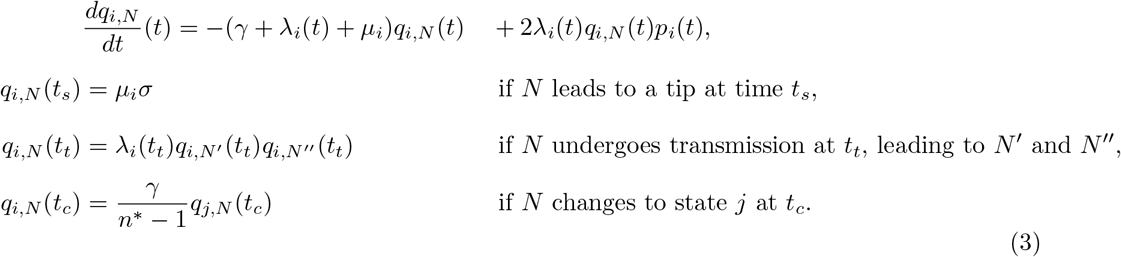

The probability of a phylogeny starting at root time τ with initial state *I* is *q*_*I*,*N*_(τ) so the full likelihood can be calculated from Eq 3. Rather than writing it recursively as in Eq 3, it can be written as a closed form equation by defining the edge likelihood function 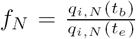 for an edge *N* in state *i* with start time *t*_*b*_ and end time *t*_*e*_. *f*_*N*_ follows the differential equation in Eq 3 with initial condition *f*_*N*_(*t*_*e*_) = 1. The full likelihood of the model *M* given the phylogeny *T* is then obtained by multiplying the likelihoods of all edges as shown in Equation (4), where *n* is the number of states (including the root state) in the tree, *N*_*i*_ is the set of edges in state *i*, *T*_*i*_ the set of transmission events in state *i* and *S*_*i*_ the set of tips in state *i.*

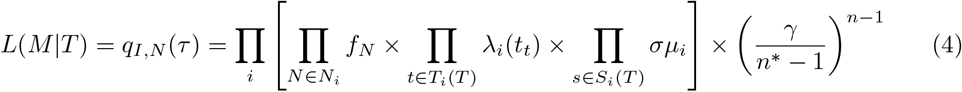

This likelihood function can be applied to trees with or without a root edge, i.e trees starting with one lineage or two at time τ.

## 2.3 Approximations to the likelihood function

### 2.3.1 Simplifying the number of states

Since the real number of clusters in the underlying network *n\** is unknown, we need to estimate it. However this parameter only appears in the likelihood in the factor 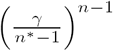 so maximizing the likelihood is equivalent to minimizing *n*.* We further assume that each migration enters a previously not visited state, i.e. *n* ≥ n.* Together, the maximum likelihood estimate will always be *n* = n.* Thus we fix *n* = n* in the inference.

### 2.3.2 Ignoring state changes in unsampled subtrees

The equations for *p* and *f*_*N*_ do not have an analytical solution. Numerical integration is computationally expensive and can be unstable for certain parameters, so we make the assumption that no state changes happen in the unsampled parts of the tree, meaning we observe all state changes in the final tree. With this assumption, the master equation for *p*_*i*_(*t*) changes to Equation (5),

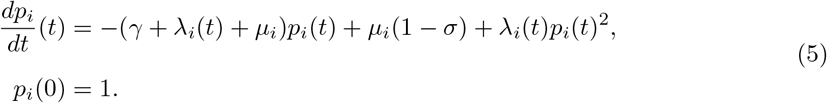

These equations have an analytical solution for constant transmission and removal rates, but not necessarily for time-dependent rates. To obtain a closed form solution, we use time discretization and assume that in each time step the transmission rate can be considered constant, as described in the next section.

### 2.3.3 Time discretization

We discretize the time-dependent transmission rates by assuming that they can be considered locally constant on small enough intervals. The grid size used for the discretization is fixed across the tree and needs to be specified by the user. A smaller size will improve the accuracy of the likelihood calculation but also increase the computational cost.

#### Time discretization for *p*

A closed form of the extinction probability and the likelihood function can be obtained for piecewise constant transmission and removal rates. Assuming constant rates in Eqn 5, and a generic initial condition *p*_*i*_(*t*_*IC*_) = *V*_*IC*_ (rather than the initial condition *p*_*i*_(0) = 1), we obtain an analytic solution of Eqn 5,

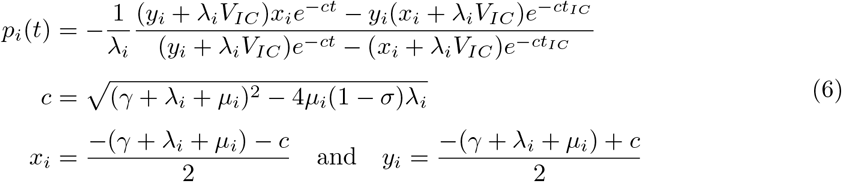

This solution can be verified by differentiating the solution and substituting the result into Eqn 5.

To obtain *p*_*i*_(*t*) using this time discretization, we divide the time interval [τ; 0] into a grid. Starting with *p*_*i*_(0) = 1, we can then evaluate *p*_*i*_ using Eq 6 in each grid interval going backwards in time, using as initial value the solution of the previous grid interval.

#### Time discretization for *f*_*N*_

A closed form solution of the edge likelihood function *f*_*N*_ can now be calculated, for a small time interval [*t*_*l*_;*t*_*l*-1_] on an edge *N* in state *i*. This expression uses the value of *p*_*i*_(*t*_*l*-1_), which can be calculated as explained in “Time discretization for *p*”. We define 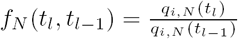, and obtain

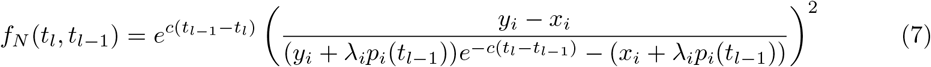

This expression for *f*_*N*_(*t*_*l*_, *t*_*l*-1_) is a solution of the differential equation 3 with *f*_*N*_(*t*_*l* -1_) = 1, assuming the rates are constant in interval [*t*_*l*_,*t*_*l* -1_] and using the approximate function *p*_*i*_(*t*) from Eq. 6. This can be easily verified by differentiating Eq. 6 and Eq. 7 and substituting the resulting expressions 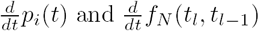 into the differential equations 5 and 3. Equations 6 and 7 are identical to the expressions used in the birth-death skyline model and a full derivation of them can be found in (17).

We now describe how to calculate *f*_*N*_ and obtain an evaluation of the likelihood provided in Eq. 4, using Eqn 7. Values of *p*_*i*_ for all branching times and state change times are precomputed to avoid the repetition of those calculations for multiple edges. For edge *N* in state *i* starting at time *t*_*b*_ and ending at time *t*_*e*_ (i.e. *t*_*b*_ < *t*_*e*_), we aim to calculate *f*_*N*_(*t*_*b*_,*t*_*e*_). Thus we aim to solve, using the time discretization, the differential equation in Eqn 3 with initial value *f*(*t*_*e*_,*t*_*e*_) = 1:

1. Fetch the precomputed value of *p*_*i*_(*t*_*e*_).
2. Divide the interval [*t*_*b*_,*t*_*e*_] in *k* equidistant intervals [*t*_*k*_, *t*_*k*-1_], [*t*_*k*-1_, *t*_*k*-2_], …, [*t*_1_, *t*_0_] with *t*_0_ = *t*_*e*_ and *t*_*k*_ = *t*_*b*_.
3. For each step *l* ∈ [1…*k*] do the following:
  a. calculate *λ*_*i,l*_ the mean of *λ*_*i*_(*t*) on the interval [*t*_*l*_,*t*_*l*-1_], then
  b. calculate *p*_*i*_(*t*_*l*_) and *f*_*N*_(*t*_*l*_,*t*_*l*-1_) by using the constant rates solutions provided in Eqn 6 for *p* and in Eqn 7 for *f* with *λ*_*i*_ = *λ*_*i,l*_, based on the value *p*_*i*_(*t*_*l*-1_) given by the precomputed value if *l* = 1 and by the previous step *l* − 1 otherwise.
4. Finally, compute 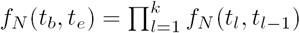.

### 2.4 Algorithm

We now present an algorithm which identifies the state change configuration and associated parameters that maximize the likelihood in Eq. 4 for a particular phylogeny *T*.

#### 2.4.1 Initial condition

The first step of the algorithm is to infer the most likely parameters for a constant rate birth-death model given the tree. These parameters will be used as starting values for the optimization in further steps. The initial values used in the optimization can have a great impact on the entire inference: if they are too distant from the optimal values, it can happen that the constant rates optimization finds only a local optima, and this will in turn affect all subsequent steps of the inference. Our method avoids this issue by applying an initial coarse-grained optimization step prior to the main optimization algorithm. Initial values are tested until no further improvement of the optima found by the optimization can be obtained. This optima will then be accepted as the global optima for the constant rates model. The user-provided starting values define the order of magnitude of the values tested in this phase.

#### 2.4.2 Maximum likelihood search

We then use a greedy approach to add state changes until no further improvement of the likelihood can be obtained. New maximum likelihood estimates are obtained for all transmission, decay, removal and state change rates each time a new state change is tested, but the positions and times of previous state changes are fixed.

Once a configuration has been found in which no more state changes can be added to improve the likelihood, we will attempt to recursively remove all the states from this configuration. This step is designed to compensate partly for the fact that the greedy approach never goes back on previous state change assignments, and so can end up in sub-optimal configurations.

Once no further improvements of the likelihood can be obtained by either adding or removing a state, the method will return the best fitting model found, including the state configuration and the maximum likelihood estimates for all parameters.

The full algorithm, including the initial coarse-grained search phase, is as follows:

1. Find the most likely parameters for a one-state birth-death model (i.e with identical birth and death rates across the tree).
2. For all edges in the tree:
  a. add a state change on this edge, then
  b. find the most likely parameters for this state configuration, then
  c. keep the edge as candidate if it is the most likely found so far.
3. If a configuration with n+1 states was found that is more likely than the configuration with n states, keep it and go back to step 2.
4. For each state change in the configuration:
  a. remove this state change.
  b. find the most likely parameters for this state configuration, and
  c. if the configuration without this state was more likely than the previous configuration, keep it.
5. If at least one state was removed, go back to step 4.
6. Otherwise, end and record the most likely model.

### 2.5 Implementation

The likelihood calculation and Maximum Likelihood inference are implemented as a publicly available R package. Partial results of the inference are automatically saved after each optimization step, so that an interrupted run can be resumed at any point. The full results returned include the best estimates for the number and positions of states, as well as all initial transmission rates, transmission decay rates and removal rates of each state. An estimation of the uncertainty around the result is provided by the maximum likelihood values found for each number of states *n* up to *ñ* + 1 where *ñ* is the maximum likelihood inferred number of states.

All analysis, pre- and post-processing of the datasets were done using custom R scripts, included in the Supplementary Materials.

#### 2.5.1 Time positions of state changes

The model and the likelihood function allow for state changes to be placed anywhere on an edge. The implementation of the algorithm allows for the time positions of changes to be estimated as additional parameters, but this is computationally expensive especially when the number of state changes grow. As a consequence we also provide the option to limit the positioning of changes to predetermined positions on edges: they can be positioned at either 10%, 50% or 90% of the length of the edge they are on. An intermediate option is also available, which will test all three predetermined options and keep the most likely.

#### 2.5.2 Speed improvement option

The algorithm as presented in the previous sections is fast at the beginning of the inference but will progressively slow down as more states are added, due to the increase in the number of parameters that need to be optimized.

We have thus added a so-called ‘fast optimization’ option, which limits the number of parameters which are allowed to change during one step of the maximum likelihood optimization. In practice, when adding the n-th state change, only the parameters *λ*_0,*n*+1_, *λ*_0,*a*_, *z*_*n*+1_, *z*_*a*_, *µ*_*n* +1_ and *µ*_*a*_ are optimized, where *a* is the state ancestral to the new state change. All other parameters are fixed to the values inferred when adding the *n*-th state. Thus this option results in each step of the algorithm having a constant cost instead of a cost dependent on *n*, however it will lose some precision by fixing parameters.

It is to be noted that it is possible to run the normal analysis for the early steps of the algorithm and turn on the fast optimization afterwards.

## 3 Results

### 3.1 Cluster inference on simulated data

#### 3.1.1 Dataset

We use a simulated dataset produced by (10). This dataset contains simulated epidemics on three different types of networks, A, B and C. The network structure A is composed of 13 communities of 20 subjects each, with each community being a fully-connected graph and one bridge linking any two communities.

The network structure B consists of one central community of size 60, representing a main sexual contact network, connected by single bridges to 25 communities of size 20. Each small community is a fully-connected graph. The set of small communities represents disjoint sexual contact subnetworks in a population of interest.

The C networks are made of 100 communities each. The size of those clusters was sampled from a distribution obtained from a phylogeny of the Swiss HIV Cohort Study (SHCS) dataset (see (10) for details). To ensure that all communities are accessible, they are first linked in a chain. Additional bridges are then created by connecting any two vertices belonging to different communities with probability 0.00075.

In all types of networks, edges between communities are weighted, with the weight value 0.25, 0.5, 0.75, or 1. This means that the rate of transmission on these edges is respectively 25%, 50%, 75 % and 100% of the transmission rate on within-community edges.

Epidemics were simulated on these networks starting from one random introduction in A networks, one random introduction in the main community in B networks, and two random introductions in C networks. All infected individuals were sampled upon removal and a transmission tree was built from the sampled tips. Thus there is no phylogenetic uncertainty in this dataset: the tree represents exactly the progress of the simulated epidemic. For each type of network (A,B,C) and each weighting scheme (w=0.25,0.5,0.75 or 1), 300 epidemics were simulated, for a total dataset of 3600 trees.

Network structure B was designed to correspond best to the monophyletic assumption of the cutpoint-based clustering methods: the epidemic starts in the main cluster and the smaller islands are not connected with each other so all infections originating from the same population cluster will be grouped in a single clade. Network structure A, on the other hand, allows for the possibility of multiple introductions in the same population cluster and nested clusters, thus breaking some of the assumptions of the cutpoint-based methods.

Various features of the A,B,C networks and the resulting simulated trees are shown in table 1. Networks A and B are very similar both in the size of their trees and in the cluster partition inside trees. Network C, on the other hand, contains a large number of fairly small clusters. Even though C trees are much larger on average, the clusters they contain are very small on average and 34% of them include only 1 or 2 tips of the tree. These very small clusters contain very little signal from the underlying contact network, and thus are not expected to be detected by the method.

**Table 1:** General features of the A, B, C networks. All numbers are averages over the 4 weighting schemes, i.e averages over all 1200 trees in each network.

| Network type | A | B | C |
| --- | --- | --- | --- |
| Number of clusters in the network | 13 | 26 | 100 |
| Number of elements per cluster | 20 | 21.54 | 9.84 |
| Number of tips per tree | 52 | 60 | 196 |
| Number of clusters per tree | 5.95 | 6.63 | 39.10 |
| Number of elements per cluster in the tree | 9.45 | 9.57 | 5.17 |
| Proportion of small clusters (<3 elements) in trees | 21% | 14% | 34% |

#### 3.1.2 Comparison with cutpoint-based methods

We ran our maximum likelihood inference on the trees and inferred clusters by considering all tips in the same state to be coming from the same community. In accordance with the simulation conditions we set *σ* = 1 in the inference. The removal rates *µ*_*i*_ are assumed independent of the population cluster, and so they are set to the same value *µ* for all states. The time positions of the state changes were fixed using the intermediate option of testing positions at 10%, 50% and 90% of the length of the edges the state changes were on.

The correspondance between the real network communities and the clusters inferred from the tree was assessed using the Adjusted Rand Index (ARI) (18; 19). We compare the results from our method to the results obtained by (10) using cutpoint-based clustering methods.

Figure 2 shows the scores obtained by our MSBD method on the simulated A,B,C networks compared to the scores of the cutpoint-based clustering methods. All methods used the same cutpoints values, except for the method based on Definition 3 (Def3). Data corresponding to this method was rescaled to fit in the same figure. As shown in (10), the results of the cutpoint-based methods are highly variable and good scores can only be obtained from a narrow range of cutpoints. In addition, the best cutpoint value is highly dependent on the underlying network structure: in methods other than Def3, the best scores are obtained for a cutpoint of *c* ≈ 0.15 for networks A, *c* ≈ 0.03 for networks B and *c* ≈ 0.02 for networks C. For Def3, the best score is obtained for *c* ≈ 0.05 for networks A, *c* ≈ 0.16 for networks B and *c* ≈ 0.04 for networks C. We define the “peak range” of cutpoints for each method, network structure and weighting scheme as the range of cutpoints which give a score which is at least 75% of the best score obtained for any cutpoint. With this definition the peak ranges are very narrow, with an average length of respectively 0.008, 0.015 and 0.016 for networks A, B and C in methods other than Def3. The peak ranges obtained with Def3 are much wider, but a direct comparison is difficult due to the different definition used for the cutpoint. In all methods the peak ranges for networks A and C on one hand, and B on the other hand have very little overlap and the best cutpoint for C is never found in the peak range of either A or B, and vice-versa. In conclusion it is impossible to get good results from all network types with any single cutpoint value.

**Figure 2:**
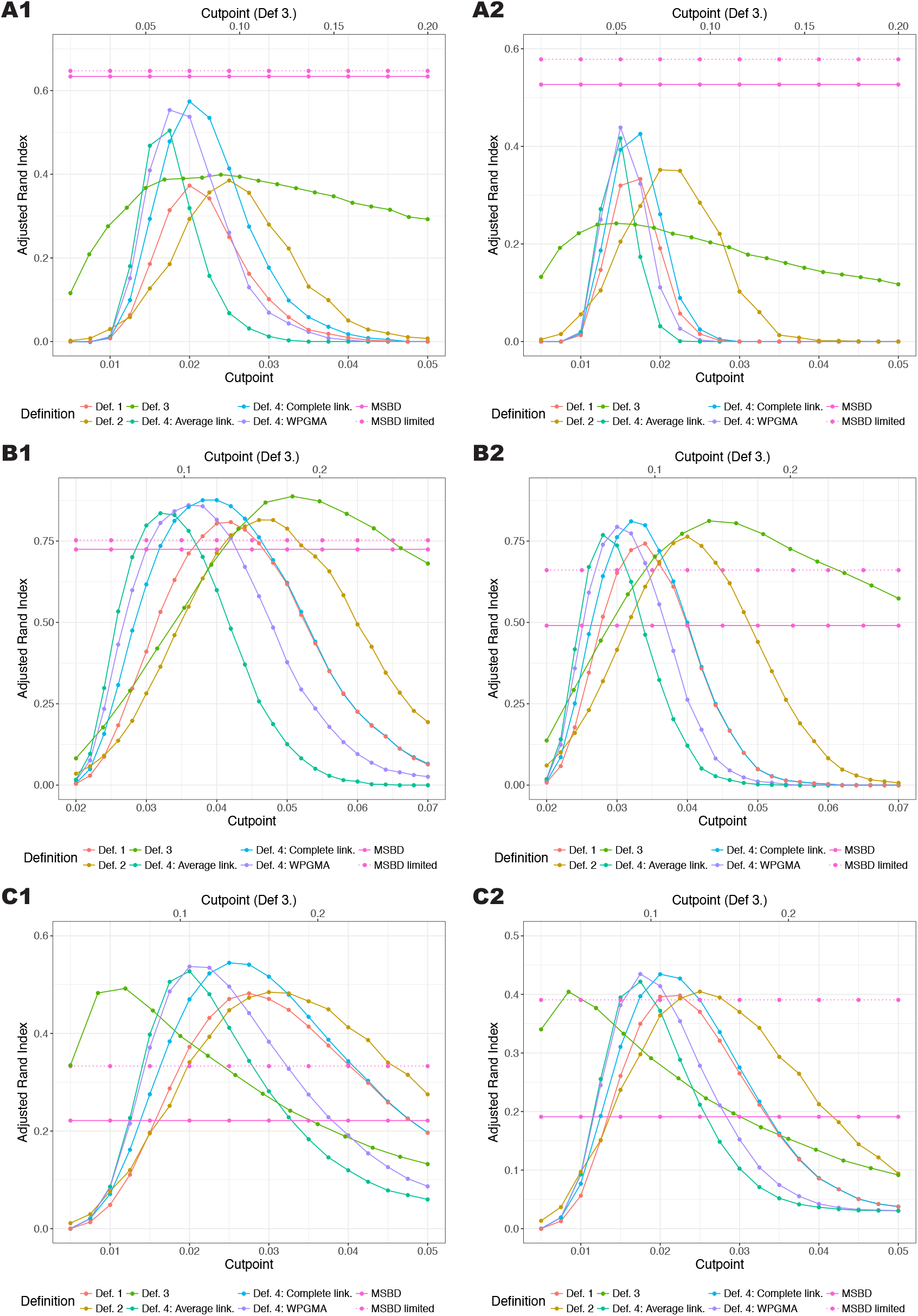
Comparison of the average ARI obtained by the different clustering methods in function of the set cutpoint on networks A (parts A1,A2), B (parts B1,B2) and C (parts C1,C2). For each network the first column (part 1) shows the results for weight *w* = 0.25 and the second column (part 2) for *w* = 1. Our proposed MSBD method is not dependent on a cutpoint. The cutpoint range for Definition 3. is shown on the x-axis on the top, the cutpoint range for all other definitions are shown on the x-axis at the bottom.

In addition the cutpoint-based methods are sensitive to network features and in particular to the non-respect of the monophyletic assumption. In both the A and C networks, the best score obtained by any cutpoint-based method is ≈ 0.45 for the weighting scheme *w* = 0.25 and ≈ 0.55 for *w* = 1, whereas it goes up to ≈ 0.85 and ≈ 0.9, respectively, in networks B.

In comparison, the MSBD method performs less well on B networks, with an average score of 0.73 for *w* = 0.25 and 0.49 for *w* = 1. However, it performs much better on A networks, with an average score of 0.64 for *w* = 0.25 and 0.53 for *w* = 1. The worst results are obtained on the C networks, where the average score is ≈ 0.2 for all weights, less than half the best scores obtained by cutpoint-based methods.

The low scores obtained on the C networks point to a potential limitation of our method on the number of clusters that can be inferred from a tree. As seen in the network features, the trees simulated on the C networks contain clusters which have less elements on average, and a higher proportion of very small clusters. These clusters may be harder to detect due to their low signal. To confirm this hypothesis we calculated the scores obtained by the MSBD method when excluding all tips that belonged to a cluster with strictly less than 8 tips. The results are shown in figure 2 (dotted line). The proportion of tips excluded by applying this criteria is shown in table 2. The scores of all network structures and all weighting schemes improved when applying this criteria. The improvement increased with the proportion of tips belonging to the excluded clusters, supporting our hypothesis that the MSBD method has difficulty identifying them. In particular, the MSBD scores on the C network structure for weight ≥ 0.5 increase to a level on par with the scores obtained by cutpoint-based methods.

**Table 2:** Proportion (%) of tips belonging to clusters with strictly less than 8 tips, per network structure and weighting scheme.

| Network type | A | B | C |
| --- | --- | --- | --- |
| $w = 0.25$ | 9.02 | 9.87 | 33.9 |
| $w = 0.5$ | 15.3 | 16 | 44.6 |
| $w = 0.75$ | 21.7 | 20.8 | 51.4 |
| $w = 1$ | 25.1 | 22.2 | 55.5 |

### 3.2 Quality of the parameter inference

To evaluate the performance of our MSBD method beyond cluster identification, we simulated several datasets of 200 trees each under the multi-state birth-death process, with various parameter combinations. Simulations were done using Gillespie’s algorithm. Tips were sampled upon removal and the process was ran until the tree reached 50 sampled tips. Since these trees were not built from network simulations we did not try to assess the quality of the cluster inference, but we focused on the quality of the parameter inference and on whether our method can adequately distinguish between trees that contain several clusters and trees that do not.

The results are summarized in Table 3. We can see that although the MSBD method is able to consistently infer multiple clusters when they are present, it will also wrongly detect one additional cluster in around 25% of the trees that only contain one cluster. This may be a problem of noise, where due to the stochasticity of the simulation one subtree is slightly more likely to be attributed different rates than the rest of the tree. This problem can be alleviated by looking at the difference in the inferred transmission rates of each cluster, which are also outputted by our method: a smaller difference is more likely to be indicative of noise. As previously noted, the method also tends to underestimate the number of clusters in multi-cluster trees, mostly because it cannot detect clusters below a certain size.

**Table 3:** Parameter inference on several datasets. Each dataset contains 200 trees of 50 tips each, simulated under a multi-state birth-death process using Gillespie’s algorithm. Transmission rates are averaged over the entire tree.

| Dataset parameters | | $\lambda_0 = 25, z = 12,$<br>$\mu = 1, \gamma = 0$ | $\lambda_0 = 25, z = 15,$<br>$\mu = 1, \gamma = 0$ | $\lambda_0 = 10, z = 1,$<br>$\mu = 5, \gamma = 0.5$ | $\lambda_0 = 10, z = 2,$<br>$\mu = 5, \gamma = 0.5$ |
| --- | --- | --- | --- | --- | --- |
| Average number of clusters | simulated | 1 | 1 | 4.95 | 6.38 |
|  | > 5 individuals, simulated | 1 | 1 | 1.92 | 2.49 |
|  | inferred | 1.22 | 1.25 | 2.43 | 2.65 |
| Average transmission rate | simulated | 1.09 | 0.86 | 6.95 | 5.40 |
|  | inferred | 1.54 | 1.38 | 7.52 | 6.20 |
|  | median absolute error | 0.37 | 0.49 | 0.75 | 0.78 |
| Average removal rate | simulated | 1.0 | 1.0 | 5.0 | 5.0 |
|  | inferred | 0.88 | 0.91 | 4.64 | 4.50 |
|  | median absolute error | 0.21 | 0.20 | 0.73 | 0.71 |

Regarding the parameter inference, the method has a slight bias towards overestimating the transmission rate and underestimating the removal rate. This is potentially due to our simulation process being conditioned on reaching 50 tips, which could bias datasets in favour of trees showing apparent higher diversification rates. Overall, the absolute error on the inferred parameters remain low compared to the true values, both in datasets with one cluster and in datasets with multiple clusters.

In conclusion, the parameter inference from the MSBD method is reliable, although it suffers from noise when applied to trees which contain only one cluster.

### 3.3 Speed improvement option

In this section we compared the performance of the “fast optimization” option and the regular algorithm. We used a dataset of 300 trees of average size 200 tips, on which a partial inference had already been performed, so the algorithm started from a saved state in which multiple state changes had already been found. One optimization step of the algorithm was then performed, i.e the inference added a state change on one edge of the tree. As shown in figure 3, we measured both the speed-up resulting from using the faster option and the difference in the maximum log likelihood found.

**Figure 3:**
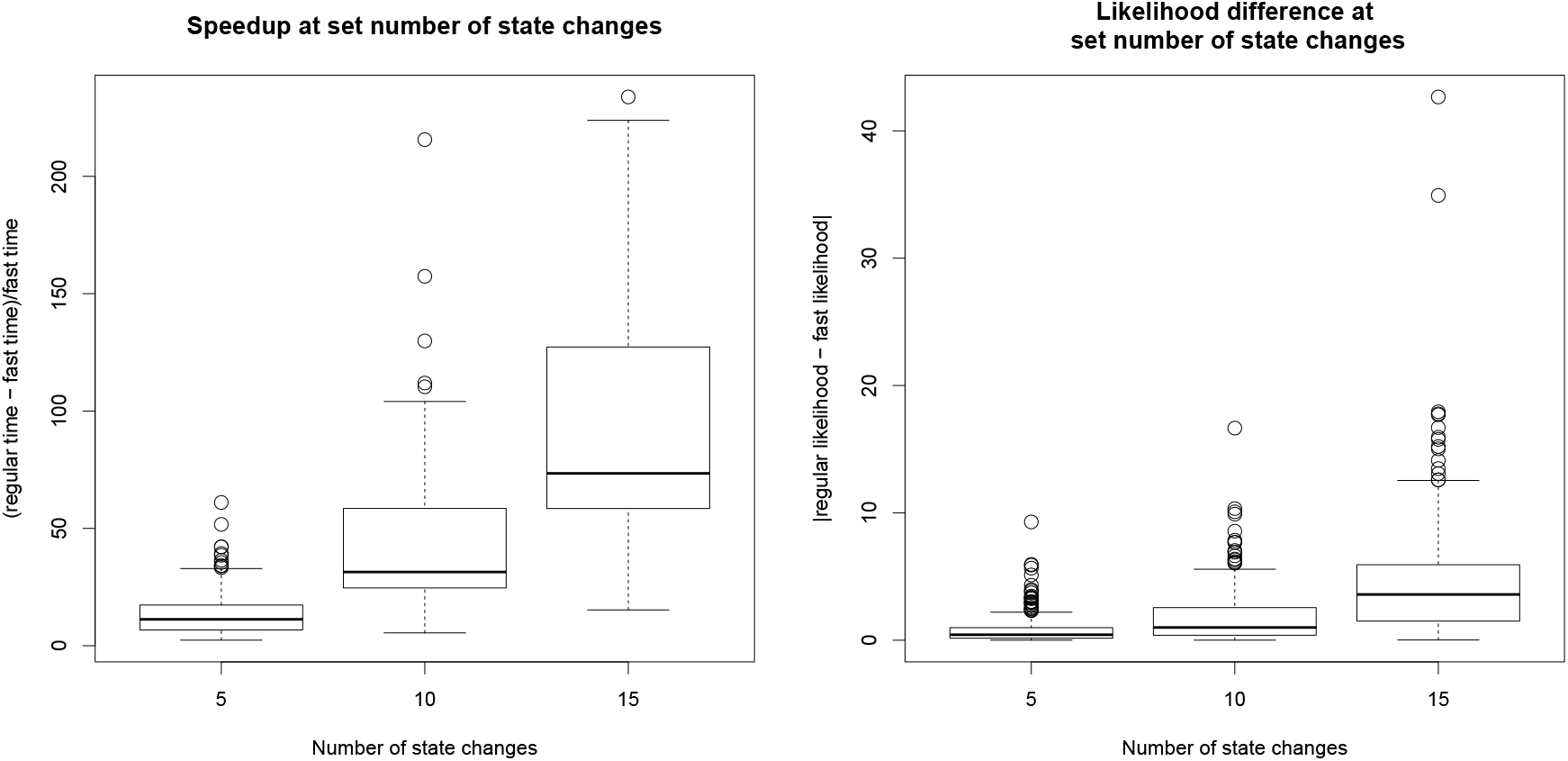
Box plots representing the speed-up (A) and likelihood difference (B) on one step of the algorithm when using the ‘fast optimization’ option compared to the default settings. The dataset used was divided in three categories based on the number of state changes already found by the inference before the test was run.

As expected the speed-up achieved increases with the number of states already present in the tested configuration. At 5 state changes, the fast optimization is on average 10 times faster than the regular one, with a number of outliers with speed-ups of up to 50 times. At 15 state changes the speed-up is of 70 on average, a considerable improvement. The difference in the maximum log-likelihood obtained using the less-precise fast option also increases with the number of state changes, although the difference remains small compared to the log-likelihood value, which is on average -1690 for the regular optimization across all categories. The runtimes for one edge are on average 170s at 5 state changes and 1250s at 15 state changes for the regular optimization. Since every step of the algorithm involves testing all edges of the tree, the “fast” option is thus necessary to ensure completion of the inference on trees with more than 10 clusters.

### 3.4 Cluster inference on HIV dataset

In this section we analyze a tree used in another study of the correlation between sexual networks and tree features, (9). HIV-1 subtype B pol sequences were obtained from the Swiss HIV Cohort Study 192 (SHCS). While the Swiss epidemic includes a mixture of population risk groups including heterosexuals, injection drug users and MSM, only viral samples from MSM were analyzed. A large cluster including 200 sampled individuals who predominantly lived or sought treatment in the Zürich area was identified from a maximum likelihood (ML) phylogeny of the complete dataset. The phylogeny of this cluster was then obtained by fitting a SIR-type pairwise epidemic model to this sub-epidemic while simultaneously inferring the tree from the sequence data in BEAST2. We re-analyze the tree provided for that cluster in the Supplement of (9), this is a random tree from the posterior sample.

The results of the MSBD analysis on cluster 581 are shown in figure 4, part A. Three sub-clusters are identified in the tree, one with a higher base transmission rate than in the backbone of the tree, and two with similar base transmission rates which are lower than in the backbone of the tree.

**Figure 4:**
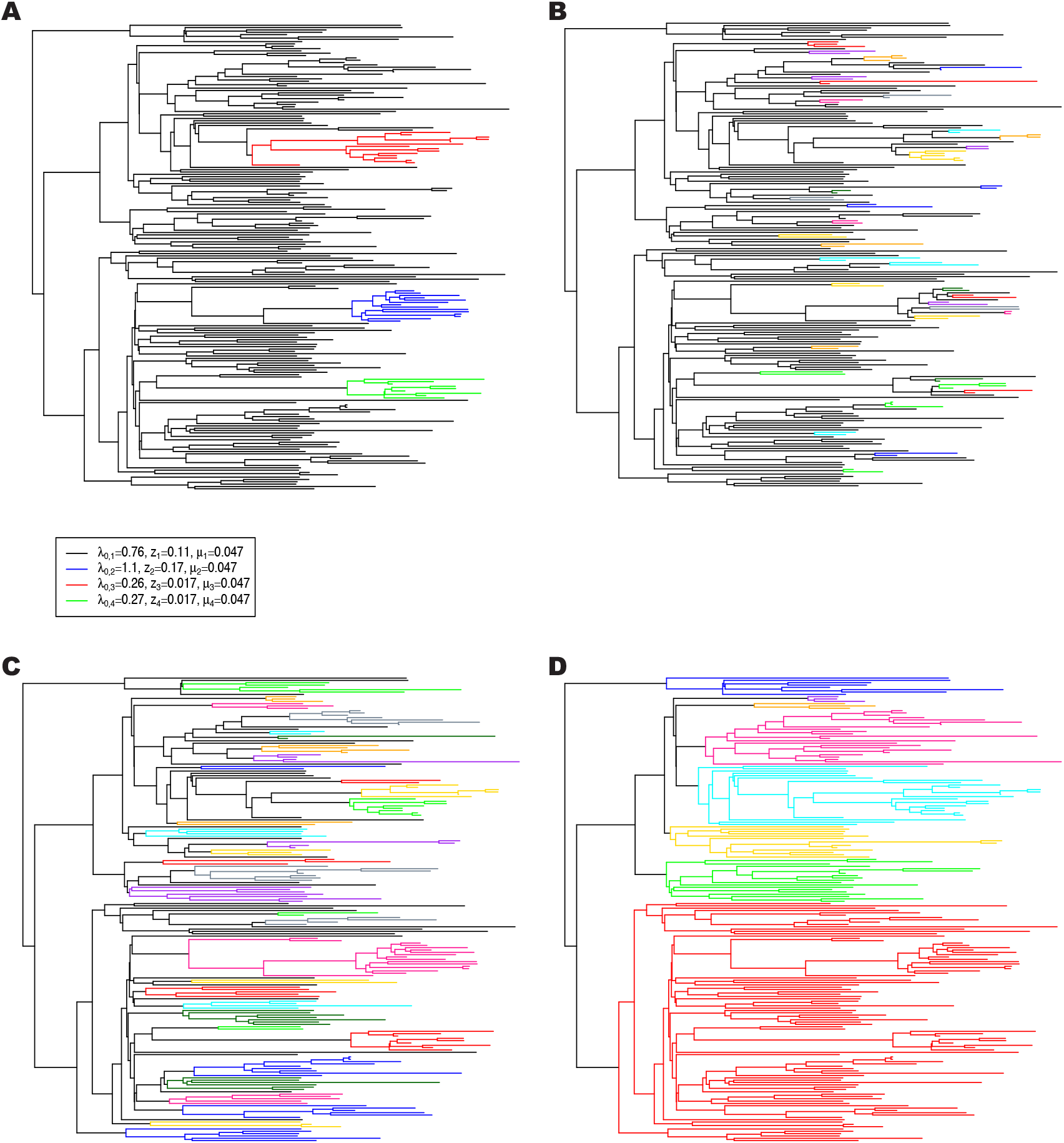
Comparison of the clusters obtained with MSBD (part A) or with Cluster Picker with a bootstrap threshold of 0.0 and a genetic distance threshold of 1.5% (part B), 4.5% (part C) and 8% (part D). Cluster Picker only identifies monophyletic clades as clusters, so each coloured clade is a separate cluster.

We compare our results to results obtained using the software Cluster Picker (20), which detects clusters based on a combination of genetic distance between tip sequences and bootstrap support at the nodes. It relies on two user-defined thresholds for both these measures, and so it is a cutpoint-based method. Genetic sequences were generated for that tree using the software SeqGen (21), using a GTR model with a gamma distribution with 4 rate categories and invariant sites. The parameters of the molecular evolution model were set to the estimates obtained by (9) when inferring the tree, which are shown in table 4.

**Table 4:** Parameter values used to simulate sequences with SeqGen.

| Parameter | Value used |
| --- | --- |
| Proportion of invariant sites | 48% |
| Frequency of A | 0.38 |
| Frequency of C | 0.16 |
| Frequency of G | 0.22 |
| Frequency of T | 0.24 |
| Shape of gamma heterogeneity | 0.57 |
| Substitution rate | 0.0015 |
| Transition rate $A \rightarrow C$ | 0.23 |
| Transition rate $A \rightarrow G$ | 1.12 |
| Transition rate $A \rightarrow T$ | 0.09 |
| Transition rate $C \rightarrow G$ | 0.14 |
| Transition rate $C \rightarrow T$ | 1.0 |
| Transition rate $G \rightarrow T$ | 0.11 |

As with other cutpoint-based methods, the results depend strongly on the user-defined values. We used three different cutpoint values for the genetic distance: 1.5%, 4.5% and 8%. 4.5% is the default value proposed by Cluster Picker and is the higher bound of the range recommended by Cluster Picker for HIV data, whereas 1.5% is the lower bound of the recommended range. For the bootstrap support threshold we used the value 0.0. With this cutpoint the bootstrap support is disregarded entirely, which mimmicks the behaviour of the methods studied by (10). The results are shown in figure 4. We can see that the number of identified clusters is strongly dependent on the cutpoint values, in keeping with the results obtained by (10). The size of the identified clusters varies also widely, even within the bounds of the recommended range of cutpoints. With the default setting of 4.5%, the clusters identified by MSBD are also recovered with Cluster Picker, although one of the clusters is split in two in the clustering by Cluster Picker.

## 4 Discussion

We have introduced a novel method of identifying transmission clusters from a phylogeny, based on a multi-state birth-death (MSBD) model. Our likelihood function makes two important assumptions: the first one is that all the clusters in the transmission network appear in the tree, and the second one is that unsampled subtrees, i.e subtrees that do not appear in the reconstructed phylogeny, do not contain new introduction events. The implementation also relies on a time discretization which approximates all transmission rates as locally constant on small time intervals. A similar discretization can be applied to extend our method to time-dependent removal rates, although the current implementation only allows removal rates to vary with state, not through time.

This new method has a few key differences compared to the cutpoint-based clustering methods. Firstly, it is not restricted to monophyletic clades and can find clusters that are nested within one another in the phylogeny. As a result our method clearly outperformed the others on trial networks which were designed specifically to violate the monophyletic assumption. Secondly, as the MSBD method is model-based it does not rely on an arbitrary cutpoint; this is particularly important as (10) found that the quality of the clusters obtained by the cutpoint-based clustering methods was extremely dependent on the value of this cutpoint, on all network types. As seen in the results it is not possible to define a single cutpoint value as adequate for all network types, which limits the usefulness of cutpoint-based methods in the absence of prior infomation on the transmission network. The chosen cutpoint value is strongly linked with the number of clusters inferred by cutpoint-based methods, thus obtaining the correct clusters requires prior knowledge on the true number of clusters. Overall, while our method may not perform as well on certain types of network as cutpoint-based methods, it is more reliable and consistent in its results and does not require additional information from the user to get optimal results.

As seen from the low scores obtained on the more fragmented trial networks and the improvements obtained by limiting the size of the clusters to be detected, the MSBD method has a strong limitation on the size of clusters that can be inferred from a tree. Contrary to the cutpoint-based methods, which can handle arbitrary numbers and sizes of clusters, our method can only add clusters when there is a strong signal for them and thus performs poorly in datasets with many small clusters. As before though it should be noted that this low performance is compared to the optimal results obtained by cutpoint-based methods, which require reliable information on the expected number of clusters.

Another limitation of the current implementation is its computational cost, which limits the size of the trees that can be analyzed in a reasonable time to a few hundred tips. Improving the speed was the reason for several approximations such as the limitations on the positions of state changes and the ‘fast optimization’ option, however these options necessarily limit the precision of the results. Future work will focus on implementing the algorithm in parallel in order to address this limitation.

Finally as a result of using a Maximum Likelihood framework, estimating the uncertainty around the various estimated parameters is problematic, in particular for the positions and number of state changes. The current implementation will estimate the uncertainty around the number of states *n* by returning the best likelihood values for each *n* tested, although we expect that better estimates could be achieved using a Bayesian framework.

## 5 Acknowledgements

We thank Dr. Timothy Vaughan for his comments and suggestions on the manuscript and Dr. Luc Villandré for his help with using the network simulation dataset.

## 7 Author contributions

JBS implemented the model, performed the simulations, analysed the data and drafted the manuscript. TS conceived of the study, designed the study, coordinated the study and helped draft the manuscript. All authors gave final approval for publication.

